# SPC-Clean: A napari Plugin for Reducing Speckle and Isolated Pixel Noise in Fluorescence Microscopy Images

**DOI:** 10.64898/2026.08.24.744862

**Authors:** Pendar Alirezazadeh, Elena Marie Kirsch, Yuan Tian, Joerg Bewersdorf, Jens Rittscher, Philipp Mergenthaler

**Affiliations:** Charité – University Medical Center Berlin, Center for Stroke Research Berlin, Berlin, Germany; Charité – University Medical Center Berlin, Department of Neurology with Experimental Neurology, Berlin, Germany; Department of Cell Biology, Yale School of Medicine, New Haven, CT, USA; Department of Biomedical Engineering, Yale University, New Haven, CT, USA; Department of Physics, Yale University, New Haven, CT, USA; Institute of Biomedical Engineering (IBME), Department of Engineering Science, University of Oxford, Oxford, United Kingdom; Big Data Institute, Li Ka Shing Centre for Health Information and Discovery, University of Oxford, Oxford, United Kingdom; Ludwig Institute for Cancer Research, Nuffield Department of Clinical Medicine, University of Oxford, Oxford, United Kingdom; Radcliffe Department of Medicine, University of Oxford, Oxford, United Kingdom

## Abstract

**Summary:** Speckle artifacts and isolated foreground pixels are common in fluorescence microscopy and can interfere with segmentation and subsequent quantitative image analysis. Conventional denoising methods often modify image intensities through filtering or smoothing, potentially altering biologically relevant fluorescence signals. We introduce Sparse Pixel Cluster Cleaning (SPC-Clean), a topology-aware method that removes poorly supported foreground pixels through iterative neighborhood analysis of a thresholded mask. SPC-Clean is deterministic, training-free, preserves original fluorescence intensities for practical microscopy workflows.

**Availability and implementation:** SPC-Clean is available as an open-source Python Napari plugin for microscopy image processing. Source code, documentation, and example data are available at: https://github.com/MergenthalerLab/SPC-Clean.git.

## Introduction

Fluorescence microscopy is widely used in biological and biomedical research to visualize cellular and subcellular structures through fluorescent labeling [Lichtman and Conchello [2005], Jonkman et al. [2020]]. However, fluorescence images are frequently affected by photon noise, detector limitations, background fluorescence, and acquisition fluctuations, which can produce isolated bright pixels and small speckle-like clusters that interfere with segmentation and quantitative analysis.

Conventional approaches, including Gaussian, median, bilateral, and non-local means filtering, reduce noise by directly modifying image intensities [Haider et al. [2016]]. Deep-learning methods such as CARE and Noise2Void provide powerful alternatives [Weigert et al. [2018], Krull et al. [2019]], but they operate directly on intensity values without explicit structural or topological constraints, which can lead to hallucinated or unsubstantiated image details [Belthangady and Royer [2019]] and, in blind-spot approaches specifically, to checkerboard artifacts or degraded fine-structure preservation [Höck et al. [2022]]. Learning-based approaches may additionally require curated training data, careful model optimization, and substantial computational resources [Belthangady and Royer [2019], Buhn et al. [2026]].

In many microscopy workflows, quantitative analysis relies on threshold-derived foreground masks rather than directly on denoised intensities [Ljosa and Carpenter [2009]]. Building on this observation, we propose Sparse Pixel Cluster Cleaning (SPC-Clean), a topology-aware method that first applies an intensity threshold to the original fluorescence image to generate a binary foreground mask. SPC-Clean then iteratively identifies and removes poorly supported foreground pixels and sparse clusters from this mask based on local neighborhood consistency. The resulting refined mask is finally applied to the original fluorescence image, removing the corresponding artifacts while preserving the original intensity values of the retained foreground regions.

SPC-Clean is deterministic, computationally lightweight, and requires no training data. For practical integration into microscopy workflows, it is provided as Python plugin for the *napari* image viewer [Sofroniew et al. [2022]].

### SPC-Clean: Neighborhood Topology-Based Noise Reduction

SPC-Clean is a deterministic, topology-aware method for removing isolated foreground artifacts and threshold-induced speckle from fluorescence microscopy images. As illustrated in Figure 1, the method consists of three stages. First, a global intensity threshold *H* is applied to the original image *I* to generate an initial binary foreground mask:

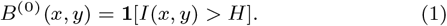

**Figure 1.**
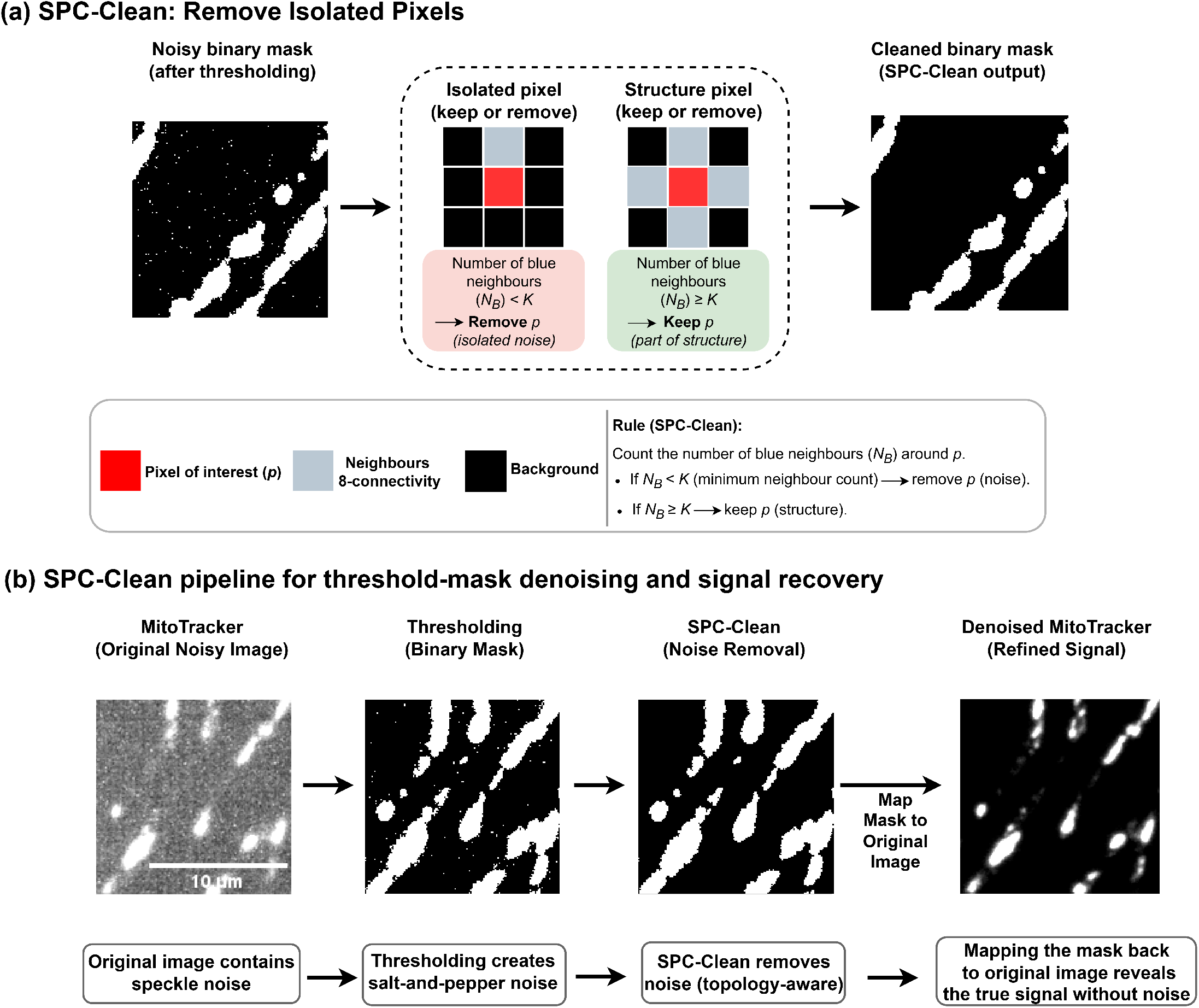
SPC-Clean workflow and neighborhood-based refinement strategy. (a) Illustration of the SPC-Clean neighborhood-consistency rule. For each foreground pixel (red), the number of active neighboring pixels (grey) within an 8-connected neighborhood is evaluated. Pixels with insufficient local neighborhood support, defined as the number of foreground pixels within their 8-connected neighborhood, are removed, whereas pixels belonging to structurally coherent regions are retained. The process is applied iteratively until convergence, progressively eliminating isolated foreground artifacts while preserving connected structures. (b) Overview of the SPC-Clean workflow, illustrated using a representative MitoTracker ExM fluorescence microscopy image. The original image is first converted into a binary mask through intensity thresholding. Although thresholding separates foreground from background, it can also produce isolated foreground pixels and speckle-like artifacts. SPC-Clean subsequently refines the thresholded mask by removing foreground pixels and isolated clusters that do not meet the required local neighborhood criterion, thereby producing a clean binary mask. Importantly, SPC-Clean does not directly modify the raw fluorescence intensity values. After thresholding, the cleaning procedure is applied only to the binary foreground mask. The original image is used again at the final reconstruction step, where the refined mask is multiplied element-wise with the original image. In this way, foreground pixels retained by SPC-Clean keep their original fluorescence intensity values while the identified foreground artifacts are removed.

Second, each foreground pixel is evaluated according to the number of active pixels in its 8-connected neighborhood. Pixels with fewer than *K* foreground neighbors are removed:

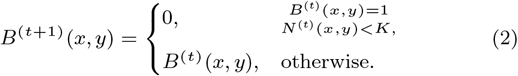

where *N* ^(*t*)^(*x, y*) denotes the local foreground support. This process is repeated until no further pixels are removed, yielding the refined mask *B*^*∗*^.

Finally, the refined mask is applied to the original image:

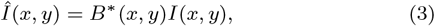

thereby removing the corresponding speckle artifacts while preserving the original fluorescence intensities of the retained foreground pixels. Importantly, SPC-Clean does not directly modify the raw fluorescence intensity values. After thresholding, the cleaning procedure is performed exclusively on the binary foreground mask. The original image is used again only at the final reconstruction step, where the refined mask is applied to retain the original fluorescence intensities of the remaining foreground pixels. Unlike intensity-domain filtering, SPC-Clean therefore suppresses artifacts through foreground-mask refinement without smoothing or modifying the intensity values of the retained signal. Detailed formulation, convergence analysis, and implementation considerations are provided in Supplementary Section S1 (Detailed Description of the SPC-Clean Algorithm).

### Computational Efficiency and Convergence

SPC-Clean is guaranteed to converge because foreground pixels are only removed and never reinstated, producing a monotonically decreasing foreground mask over a finite image domain. Each iteration has linear complexity, *O*(|Ω|), and the overall complexity is *O*(*T* |Ω|) for *T* iterations. For the parameter setting used throughout this study (*H* = 128, *K* = 3), convergence was reached after 2.92 *±* 2.17 iterations on average (range: 1-10) across 334 fluorescence image planes, demonstrating that only a few pruning passes are typically required. A detailed convergence analysis across different values of *H* and *K* is provided in Supplementary Section S2 and Supplementary Table 1.

## Results

SPC-Clean was quantitatively evaluated on three independent fluorescence microscopy acquisitions of MitoTracker-stained primary human fibroblasts subjected to pan-Expansion Microscopy (pan-ExM) [M’Saad and Bewersdorf [2020]], comprising 1,002 images in total. Detailed dataset characteristics and evaluation procedures are provided in Supplementary Section S3. Per-dataset and pooled results are reported in Supplementary Section S4 (Experimental Results), Tables 2 and 3.

Across the three datasets, SPC-Clean consistently improved foreground–background separation. The most pronounced effect of SPC-Clean was the reduction of small foreground components, which directly reflects the type of artifact targeted by the method. Pooled speckle count decreased by 96.9%, from 1113 *±* 1118 to 35 *±* 73 components per image, while speckle fraction decreased by 70.2%, from 0.976 *±* 0.029 to 0.291 *±* 0.308. This effect was consistently observed across the three independent acquisitions, with reductions in speckle count of 99.0%, 95.9%, and 97.4% for Runs 1–3, respectively. Importantly, the pronounced reduction in small foreground components was achieved while maintaining very high structural similarity of the intensity-weighted foreground signal. Structural similarity index measure (SSIM) was computed between the threshold-masked raw image and the SPC-Clean-masked raw image within the bounding box containing the union of their foreground regions. The pooled SSIM was 0.992 *±* 0.005, with Run 1 reaching 0.995 *±* 0.001, corresponding to values close to the maximum SSIM of 1. This near-unity structural similarity, despite the 96.9% reduction in speckle count, provides strong quantitative evidence that SPC-Clean selectively removes small, locally unsupported foreground components while introducing only minimal alteration to the overall spatial and intensity organization of the threshold-defined fluorescence signal. Thus, the substantial suppression of targeted foreground artifacts is achieved without broad disruption of the dominant fluorescence structures.

The utility of SPC-Clean beyond the primary dataset was further examined using MitoTracker-stained HeLa cells, antibody-labeled HeLa cells, and antibody-labeled mouse kidney tissue acquired using pan-Expansion Microscopy [M’Saad and Bewersdorf [2020], Tian et al. [2025]]. Across these distinct biological and labeling conditions, SPC-Clean removed isolated foreground artifacts while retaining coherent fluorescent structures. Representative examples are provided in Supplementary Section S5 (Additional Experimental Results), Figures S5-S7.

## Discussion

SPC-Clean provides a lightweight and practical solution for refining threshold-derived foreground masks in fluorescence microscopy. Its main distinction is that artifact removal is performed on the binary foreground topology rather than by filtering the original fluorescence intensities. The refined mask is subsequently applied to the original image, such that intensity values of retained foreground pixels remain unchanged. This is particularly useful in quantitative microscopy workflows where intensity measurements should be preserved while isolated foreground artifacts are excluded.

Evaluation across three independent MitoTracker acquisitions comprising 1,002 images demonstrated a pronounced and consistent reduction of the foreground artifacts specifically targeted by SPC-Clean. The method reduced small speckle components by 96.9%, demonstrating its strong ability to suppress isolated and locally unsupported foreground components. Importantly, this extensive speckle reduction was achieved while maintaining very high structural similarity of the intensity-weighted foreground signal, with a pooled structural similarity index measure (SSIM) of 0.992 *±* 0.005, close to the maximum value of 1. The combination of a 96.9% reduction in speckle components and near-unity SSIM provides strong quantitative evidence that SPC-Clean selectively removes the targeted small foreground artifacts while introducing only minimal alteration to the overall spatial and intensity organization of the threshold-defined fluorescence signal. This balance between substantial artifact suppression and high structural similarity represents a central advantage of SPC-Clean. Qualitative evaluation on additional HeLa-cell and mouse-kidney fluorescence datasets further demonstrated the applicability of the method across different biological samples, including tissue sections, as well as different fluorescence-labeling strategies and sample-preparation conditions.

From an application perspective, SPC-Clean requires no training data or model optimization and uses only two interpretable parameters: the intensity threshold *H* and minimum neighborhood support *K*. Its deterministic and computationally lightweight formulation makes it suitable for routine microscopy analysis and batch processing. Distribution as Python plugin for *napari* further enables SPC-Clean to be incorporated directly into existing image-analysis workflows without requiring users to implement the algorithm independently.

SPC-Clean is intended specifically as a post-thresholding refinement tool rather than a general-purpose image-restoration method. Its performance consequently depends on the quality of the initial foreground mask, and genuine structures with very limited spatial support may be removed if the selected neighborhood criterion is too restrictive. Conversely, spatially coherent background artifacts may remain because they satisfy the local support criterion. Appropriate selection of *H* and *K* should therefore consider the morphology and spatial scale of the structures being analyzed. Practical parameter-selection guidelines, supported by experimental evaluation across different parameter settings, are provided in Supplementary Section S4.

Overall, SPC-Clean provides an accessible and reproducible tool for reducing isolated foreground artifacts while leaving the fluorescence intensity values of the retained foreground signal unchanged. This distinction is particularly relevant for quantitative fluorescence analysis because retained pixels preserve their original measured intensities. Its integration into *napari*, support for batch processing, and training-free operation make it particularly suited to practical fluorescence microscopy workflows. Future developments may include three-dimensional neighborhood processing and automated parameter selection to further extend the plugin to diverse imaging conditions and biological structures.

## Data and code availability

Sample image data that allow rapid evaluation of SPC-Clean were deposited on the open-science platform Zenodo (DOI: 10.5281/zenodo.21934521). The source code is openly available on GitHub (https://github.com/MergenthalerLab/SPC-Clean) and is also archived on Zenodo (https://doi.org/10.5281/zenodo.21935012).

## Acknowledgements

This work was supported by the Core Unit pluripotent Stem Cells and Organoids (CUSCO) of the Berlin Institute of Health (BIH) at Charité – University Medical Center Berlin.

## Funding

This work was supported by the Stiftung Charité to P.M. and J.R. [StC-VF-2024-59], the Berlin-Oxford Research Partnership through the Early Career Researchers program, and in part by Wellcome Trust Collaborative awards [095927, 203285] to J.R., the Einstein Foundation Berlin [EJF-2020-602, EVF-2021-619, EVF-2021-619-2, EVF-BUA-2022-694] to P.M., the Volkswagen Stiftung [9A866] to P.M.. J.R. is Visiting Fellow at Charité funded by the Stiftung Charité. P.M. is Einstein Junior Fellow funded by the Einstein Foundation Berlin.

## Conflicts of interest

J.B. is a co-founder of panluminate Inc. which develops pan-Expansion Microscopy related products. J.R. is a co-founder of Ground Truth Labs and receives funding from Novo Nordisk.

## Notes

https://github.com/MergenthalerLab/SPC-Clean

https://doi.org/10.5281/zenodo.21934521

