## Supplemental Information for "SPC-Clean: A napari Plugin for Reducing Speckle and Isolated Pixel Noise in Fluorescence Microscopy Images"

#### Supplementary Material

S1. Detailed Description of the SPC-Clean Algorithm
S2. Computational Complexity and Convergence Analysis
S3. Dataset Description and Evaluation Protocol
S4. Experimental Results
S5. Additional Experimental Results

#### S1. Detailed Description of the SPC-Clean Algorithm

Sparse Pixel Cluster Cleaning (SPC-Clean) is a deterministic, topology-aware refinement method designed to remove poorly supported foreground pixels and sparse clusters from threshold-derived fluorescence microscopy image masks. Rather than filtering on the original fluorescence intensities, SPC-Clean operates exclusively on the binary foreground topology and subsequently applies the refined mask to the original image. The complete procedure is summarized in Supplementary Algorithm 1.

**Supplementary Algorithm 1** SPC-Clean: Iterative Neighborhood Noise Pruning

**Require:** Fluorescence image $I$, intensity threshold $H$, minimum neighborhood support $K$
**Ensure:** Refined binary mask $B^*$, speckle-reduced image $\hat{I}$

*% Stage 1: Initial foreground-mask generation*

Generate the initial binary mask:

$$B^{(0)}(x, y) = \mathbf{1}[I(x, y) > H]$$

$t \leftarrow 0$

*% Stage 2: Iterative neighborhood-consistency pruning*

**repeat**

Compute foreground support for all pixels:

$$N^{(t)}(x, y) = \sum_{(i,j)\in\mathcal{N}(x,y)} B^{(t)}(i, j)$$

Update the foreground mask synchronously:

$$B^{(t+1)}(x, y) = \begin{cases} 0, & B^{(t)}(x, y) = 1 \text{ and } N^{(t)}(x, y) < K, \\ B^{(t)}(x, y), & \text{otherwise.} \end{cases}$$

$t \leftarrow t + 1$

**until** $B^{(t)} = B^{(t-1)}$

*% Stage 3: Mask-guided reconstruction*

$B^* \leftarrow B^{(t)}$

Apply the refined mask to the original image:

$$\hat{I}(x, y) = B^*(x, y)\, I(x, y)$$

**return** $B^*, \hat{I}$

##### S1.1 Mathematical Formulation

Let $I : \Omega \to \mathbb{R}$ denote a fluorescence microscopy image defined on the finite discrete pixel domain $\Omega \subset \mathbb{Z}^2$. SPC-Clean consists of three sequential operations: initial foreground-mask generation, iterative neighborhood-consistency pruning, and mask-guided reconstruction.

**Stage 1: Initial foreground-mask generation.** A global intensity threshold $H$ is applied once to the original fluorescence image to generate the initial binary mask:

$$B^{(0)}(x,y) = \begin{cases} 1, & I(x,y) > H, \\ 0, & \text{otherwise.} \end{cases} \tag{1}$$

Pixels above $H$ are assigned to the foreground. The resulting mask may contain both biologically relevant structures and spurious foreground pixels arising from background fluctuations, detector noise, or other acquisition-related artifacts. After this initialization, SPC-Clean operates exclusively on the binary mask and does not directly modify the original image $I$. The original fluorescence intensity values are therefore preserved for all foreground pixels retained by the final refined mask.

**Stage 2: Iterative neighborhood-consistency pruning.** For each active foreground pixel $(x,y)$, local spatial support is evaluated within its 8-connected neighborhood. Defining

$$\mathcal{N}(x,y) = \{(x+\Delta x,\ y+\Delta y)\ :\ \Delta x, \Delta y \in \{-1,0,1\},\ (\Delta x, \Delta y) \neq (0,0)\}. \tag{2}$$

the foreground support at iteration $t$ is

$$N^{(t)}(x,y) = \sum_{(i,j)\in\mathcal{N}(x,y)} B^{(t)}(i,j), \qquad 0 \leq N^{(t)}(x,y) \leq 8. \tag{3}$$

A foreground pixel is retained only when its local support satisfies the minimum neighborhood criterion $K$:

$$B^{(t+1)}(x,y) = \begin{cases} 0, & B^{(t)}(x,y) = 1 \text{ and } N^{(t)}(x,y) < K, \\ B^{(t)}(x,y), & \text{otherwise.} \end{cases} \tag{4}$$

The update is performed synchronously across the mask. Thus, neighborhood support at iteration $t$ is calculated from $B^{(t)}$, and all resulting removals are incorporated into $B^{(t+1)}$. Pixels removed during an iteration are never reinstated.

This iterative operation progressively removes foreground pixels that cannot maintain sufficient spatial support as the surrounding mask evolves. Consequently, an initially supported pixel may become unsupported after neighboring pixels are removed and can therefore be eliminated in a subsequent iteration. This iterative propagation distinguishes SPC-Clean from a single-pass neighborhood filter.

**Stage 3: Mask-guided reconstruction.** Iteration terminates when the foreground topology no longer changes:

$$B^{(t+1)} = B^{(t)}. \tag{5}$$

The converged mask is denoted by

$$B^* = B^{(t)}. \tag{6}$$

The final speckle-reduced fluorescence image is then obtained by applying $B^*$ element-wise to the original image:

$$\hat{I}(x,y) = B^*(x,y)\, I(x,y). \tag{7}$$

SPC-Clean therefore produces two outputs: the refined binary foreground mask $B^*$ and the corresponding intensity image $\hat{I}$. For every retained foreground pixel,

$$B^*(x,y) = 1 \quad \Longrightarrow \quad \hat{I}(x,y) = I(x,y), \tag{8}$$

demonstrating that the original fluorescence value is exactly preserved. Conversely, pixels removed from the foreground are set to zero in $\hat{I}$. No smoothing, interpolation, or modification of retained intensity values is performed.

#### S1.2 Role of the Parameters $H$ and $K$

SPC-Clean is controlled by two interpretable parameters. The intensity threshold $H$ determines the initial foreground topology and is applied only during mask initialization. The neighborhood-support parameter $K$ determines the minimum number of active neighbors required for a foreground pixel to remain in the mask.

Lower values of $K$ impose a less restrictive spatial-support criterion and therefore favor preservation of thin or sparsely connected structures. Increasing $K$ requires stronger local support and produces progressively more aggressive removal of sparse foreground regions. In this study, $K = 3$ was used because it provided an effective balance between removal of isolated foreground artifacts and preservation of mitochondrial morphology across the evaluated fluorescence microscopy datasets.

#### S1.3 Convergence

Finite convergence follows directly from the removal-only update rule. For every iteration,

$$B^{(t+1)}(x,y) \leq B^{(t)}(x,y), \qquad \forall (x,y) \in \Omega, \tag{9}$$

and therefore the corresponding foreground sets satisfy

$$\mathcal{F}^{(t+1)} \subseteq \mathcal{F}^{(t)}. \tag{10}$$

where

$$\mathcal{F}^{(t)} = \left\{ (x,y) \in \Omega \ : \ B^{(t)}(x,y) = 1 \right\}. \tag{11}$$

Thus, $\mathcal{F}^{(t)}$ forms a monotonic non-increasing sequence of subsets of the finite domain $\Omega$. Whenever convergence has not yet been reached, at least one foreground pixel must be removed:

$$\left| \mathcal{F}^{(t+1)} \right| < \left| \mathcal{F}^{(t)} \right|. \tag{12}$$

Because the initial foreground contains only a finite number of pixels, an infinite sequence of strict removals is impossible. Hence, there exists a finite iteration $T$ such that

$$B^{(T+1)} = B^{(T)} = B^{*}. \tag{13}$$

SPC-Clean therefore always reaches a fixed point. Empirically, convergence is achieved within a small number of iterations for the fluorescence microscopy images evaluated in this study; the corresponding convergence analysis is reported in Supplementary Table S1.

#### S1.4 Computational Complexity

For an image containing $|\Omega|$ pixels, neighborhood support is computed over a fixed $3 \times 3$ local window. Since the neighborhood size is constant and independent of image dimensions, each complete iteration has computational complexity

$$\mathcal{O}(|\Omega|). \tag{14}$$

If convergence requires $T$ iterations, the total computational complexity is

$$\mathcal{O}(T|\Omega|). \tag{15}$$

Memory requirements are likewise linear in the image size because only the image and binary mask representations required for the iterative update need to be maintained. Since $T$ is small in practice, SPC-Clean introduces limited computational overhead and is suitable for processing large microscopy image collections.

#### S1.5 Methodological Scope

SPC-Clean should be interpreted as a topology-based post-thresholding refinement method rather than a general-purpose intensity denoising or image-restoration technique. It does not estimate an underlying noise distribution, reconstruct missing fluorescence signal, or modify the intensity distribution of retained pixels. Instead, it evaluates whether foreground pixels identified by thresholding possess sufficient local spatial support to remain part of the foreground topology.

This distinction is particularly relevant for quantitative fluorescence analysis because retained pixels preserve their original measured intensities. SPC-Clean therefore separates the decision of whether a pixel belongs to the retained foreground from the fluorescence value associated with that pixel.

#### S1.6 Implementation and Reproducibility

SPC-Clean is implemented in Python and distributed as a plugin for the *napari* image viewer [Sofroniew et al. [2022]]. The implementation supports both interactive inspection and batch processing of fluorescence microscopy images. Because the algorithm requires only the intensity threshold $H$ and neighborhood-support threshold $K$, it does not require training data, learned parameters, or model optimization.

The binary-mask formulation and deterministic update rule ensure that identical inputs and parameter settings produce identical outputs, facilitating reproducible application across microscopy datasets. The implementation further enables integration of SPC-Clean as a lightweight post-thresholding refinement step within existing fluorescence microscopy analysis pipelines.

### S2. Computational Complexity and Convergence Analysis

#### Theoretical convergence guarantee.

SPC-Clean is guaranteed to converge in a finite number of iterations by construction. Let $B^{(t)}$ denote the binary foreground mask after the $t$-th pruning iteration and define the foreground set

$$F^{(t)} = \{(x, y) \in \Omega : B^{(t)}(x, y) = 1\}, \tag{16}$$

where $\Omega$ denotes the image domain. During each iteration, SPC-Clean evaluates the local neighborhood support of every foreground pixel and removes those that do not satisfy the minimum neighborhood-support criterion $K$. Importantly, pixels removed during an iteration are never reinstated in subsequent iterations. Therefore, the foreground sets satisfy

$$F^{(t+1)} \subseteq F^{(t)}, \tag{17}$$

which implies that the sequence of foreground masks forms a monotonic, non-increasing sequence.

Since the image domain $\Omega$ contains a finite number of pixels, the cardinality of the foreground set satisfies

$$0 \leq |F^{(t)}| \leq |\Omega|, \tag{18}$$

and decreases monotonically until no further pixels satisfy the removal criterion. Consequently, there exists a finite iteration index $T$ such that

$$F^{(T+1)} = F^{(T)}, \tag{19}$$

at which point the algorithm has reached a fixed point and terminates. Therefore, SPC-Clean is guaranteed to converge after a finite number of iterations.

#### Computational complexity.

During each iteration, the algorithm evaluates the local support of every pixel using a fixed-size $3 \times 3$ neighborhood. Since the neighborhood size is constant and independent of image dimensions, the computational complexity of a single iteration is linear in the number of image pixels,

$$\mathcal{O}(|\Omega|), \tag{20}$$

where $|\Omega|$ denotes the total number of pixels in the image. If convergence is reached after $T$ iterations, the overall computational complexity becomes

$$\mathcal{O}(T|\Omega|). \tag{21}$$

Although the theoretical worst-case value of $T$ depends on the topology of the foreground mask and on the neighborhood-support parameter $K$, the number of iterations is independent of image resolution. Consequently, provided that $T$ remains small in practice, SPC-Clean exhibits effective linear-time scaling behavior.

#### Interpretation of the number of iterations.

The number of iterations required for convergence can be interpreted as the number of successive topological pruning passes needed to remove foreground structures that do not satisfy the neighborhood-support criterion. Foreground pixels removed during the first iterations typically correspond to isolated speckles and weakly connected artifacts, whereas later iterations operate on progressively smaller residual foreground structures. Therefore, the total number of iterations depends on both the topology of the thresholded foreground mask and the selected parameter values.

In general, lower threshold values preserve a larger number of foreground pixels during initialization, thereby increasing the number of topological configurations that require iterative pruning. Similarly, larger values of the minimum neighborhood-support parameter $K$ impose stronger local consistency constraints, resulting in additional pruning passes before convergence is achieved.

#### Empirical convergence analysis.

To characterize the practical convergence behavior of SPC-Clean, we measured the number of pruning iterations required to reach convergence under different parameter settings. The analysis was performed on a MitoTracker fluorescence expansion microscopy acquisition of primary human fibroblasts imaged on an Opera Phenix spinning-disk confocal microscope (63× water objective, NA = 1.15), consisting of a 334-plane z-stack acquired with an axial step size of 0.3 µm.

For each image plane, SPC-Clean was iteratively applied until no additional foreground pixels satisfied the neighborhood-removal criterion. The number of iterations required to reach convergence was recorded for different combinations of the intensity threshold ($H$) and minimum neighborhood support parameter ($K$).

Supplementary Table 1 summarizes the observed convergence statistics.

**Supplementary Table 1** Empirical convergence statistics of SPC-Clean measured on a 334-plane MitoTracker fluorescence microscopy z-stack for different parameter configurations.

| Threshold ($H$) | Minimum support ($K$) | Iterations (mean ± SD) | Min. | Max. | # images |
|---|---|---|---|---|---|
| 128 | 2 | $2.07 \pm 1.10$ | 1 | 6 | 334 |
| 128 | 3 | $2.92 \pm 2.17$ | 1 | 10 | 334 |
| 120 | 3 | $5.30 \pm 2.16$ | 3 | 17 | 334 |
| 128 | 4 | $7.29 \pm 7.55$ | 1 | 33 | 334 |
| 120 | 4 | $9.90 \pm 8.11$ | 2 | 44 | 334 |

Across all tested parameter combinations, SPC-Clean converged after a finite number of iterations, experimentally confirming the theoretical convergence guarantee.

As expected, the number of iterations increased when either the threshold produced a denser initial foreground mask or the neighborhood-support criterion became more restrictive. Specifically, decreasing the threshold from $H = 128$ to $H = 120$ approximately doubled the average number of required iterations, while increasing the neighborhood-support parameter from $K = 2$ to $K = 4$ resulted in substantially longer pruning sequences.

For the parameter configuration used throughout this study ($H = 128$, $K = 3$), SPC-Clean converged after $2.92 \pm 2.17$ iterations on average, with a maximum of only 10 iterations across all 334 image planes. Even under considerably more restrictive parameter settings ($H = 120$, $K = 4$), convergence remained bounded, requiring at most 44 iterations.

These findings demonstrate that the number of iterations required for convergence remains small in practice and experimentally support the effective linear-time behavior of SPC-Clean discussed in the main manuscript.

### S3. Dataset Description and Evaluation Protocol

#### S3.1 Primary MitoTracker Fluorescence Dataset

The primary evaluation comprised three independent fluorescence microscopy acquisitions of primary human fibroblasts subjected to Expansion Microscopy (pan-ExM) [M'Saad and Bewersdorf [2020]]. Primary human fibroblasts were cultured on Collagen G (Sigma) coated sterile culture ware in humidified, hypoxic growth conditions (37 °C, 5 % $O_2$, 5 % $CO_2$) in Fibroblast medium (DMEM High Glucose (Gibco), 10% foetal bovine serum (FBS, Panbiotech), 1x Anti-Anti (Gibco), 1x NEAA (Gibco), 10 ng $mL^{-1}$ FGF (Peprotech), 50 µg $mL^{-1}$ Uridin (Sigma). Coverslips were coated by incubating coating medium (KO-DMEM/F12 (Gibco), 5 % FBS (Panbiotech), 1x Anti-Anti (Gibco), 100 mM Hepes (Gibco), 0.04 % CollagenG (Sigma)) for 1–3h at 37 °C followed by two consecutive washes with 1x Dulbecco's phosphate-buffered saline (DPBS) with Calcium and Magnesium (Gibco). For sample generation, 25 000 cells were seeded on 12 mm collagen G coated glass coverslips in 24 wells and grown to confluency. Before fixation, cells were stained with 500 nM MitoTracker Orange CMTMRos (Thermo Fisher) for 45min at 37 °C. Afterwards cells were subjected to fixation in 3 % formaldehyde (Electron Microscopy Science) and 0.1 % glutaraldehyde (Electron Microscopy Science) in 1x DPBS (Gibco) for 15 min at room temperature followed by expansion as described previously [M'Saad and Bewersdorf [2020]]. Samples were imaged using the Opera Phenix spinning-disk confocal microscope (Revvity) with Harmony v5.2 software. Cells were imaged with a 63x water objective (NA=1.15) using a 561 nm laser (200ms, 100 %) and 570–630 nm emission filter. Z-stacks of 334 planes were acquired with an axial step size of 0.3 µm, corresponding to a total stack height of 100 µm.

#### S3.2 Additional Fluorescence Microscopy Datasets

To examine the utility of SPC-Clean for other data sets, three additional independent fluorescence microscopy datasets acquired under different biological and labeling conditions were evaluated. These comprised MitoTracker Orange-stained HeLa cells, antibody-labeled HeLa cells, and antibody-labeled mouse kidney tissue sections. The datasets were acquired using an Andor Dragonfly 600 spinning-disk confocal microscope as described [Tian et al. [2025]].

These additional datasets were used to qualitatively assess SPC-Clean across different cell types, labeling strategies, and tissue contexts. Representative examples are presented in Supplementary Figures S5-S7.

#### S3.3 Mask Generation and Quantitative Evaluation

For each image in the primary dataset, an initial foreground mask was generated from the raw fluorescence image using intensity thresholding. This threshold-derived mask represents the pre-cleaning condition, whereas the corresponding mask obtained after applying SPC-Clean represents the post-cleaning condition. Quantitative evaluation was performed by comparing these pre- and post-cleaning conditions.

The evaluation focused on two complementary aspects of SPC-Clean performance: speckle reduction and structural preservation. Reference-free measures were used because pixel-level ground-truth labels for fluorescence speckle artifacts are difficult to define, particularly when weak biological signal and acquisition-related artifacts overlap in intensity Haider et al. [2016], Krull et al. [2019].

Speckle reduction was quantified using connected-component analysis. A speckle component was defined as a connected foreground region with an area $\leq 10$ pixels. Speckle count represents the number of such components, whereas speckle fraction represents the proportion of speckle components relative to the total number of foreground connected components. Lower values therefore indicate greater reduction of small foreground artifacts.

Structural preservation was evaluated using the structural similarity index measure (SSIM) Wang et al. [2004]. For each image, the raw fluorescence image was multiplied by the corresponding pre-cleaning and post-cleaning masks to generate two intensity-weighted foreground images. SSIM was then computed between these images within the bounding box containing the union of their foreground regions. This analysis quantifies the extent to which the spatial and intensity organization of the threshold-defined foreground signal is maintained after SPC-Clean refinement, with values approaching 1 indicating high structural similarity.

Per-dataset and pooled quantitative results are reported in Supplementary Tables 2 and 3.

#### S4. Experimental Results

Classical denoising techniques are frequently applied directly to raw fluorescence intensity images prior to segmentation in an attempt to suppress speckle noise and background artifacts. As illustrated in Figure S1, we compared five representative approaches spanning general-purpose and speckle-specific categories: Wavelet-Bayes, bilateral filtering, non-local means (NLM), the Lee filter, and the Kuan filter.

While these methods differ in their underlying assumptions, a common limitation emerges across all of them when applied to MitoTracker fluorescence data. General-purpose filters such as bilateral and NLM operate by averaging pixel intensities within adaptive neighborhoods, which effectively smooths intensity discontinuities but simultaneously blurs the sharp boundaries of mitochondrial structures and attenuates thin, low-intensity filaments that fall below the local averaging scale. Speckle-specific filters, including Lee and Kuan, are designed under a multiplicative noise model originally developed for synthetic aperture radar (SAR) imagery and assume a homogeneous signal distribution within local windows. When applied to fluorescence images, where the foreground is inherently sparse and heterogeneous, these filters fail to distinguish isolated noise pixels from genuine low-intensity mitochondrial signal, resulting in suppression of both. As visible in the magnified regions of Figure S1(c,e,f), bilateral, Lee, and Kuan filtering not only leaves residual speckle artifacts but also degrades structural boundaries and reduces the intensity of real foreground regions, introducing distortions that propagate directly into downstream segmentation and morphological quantification.

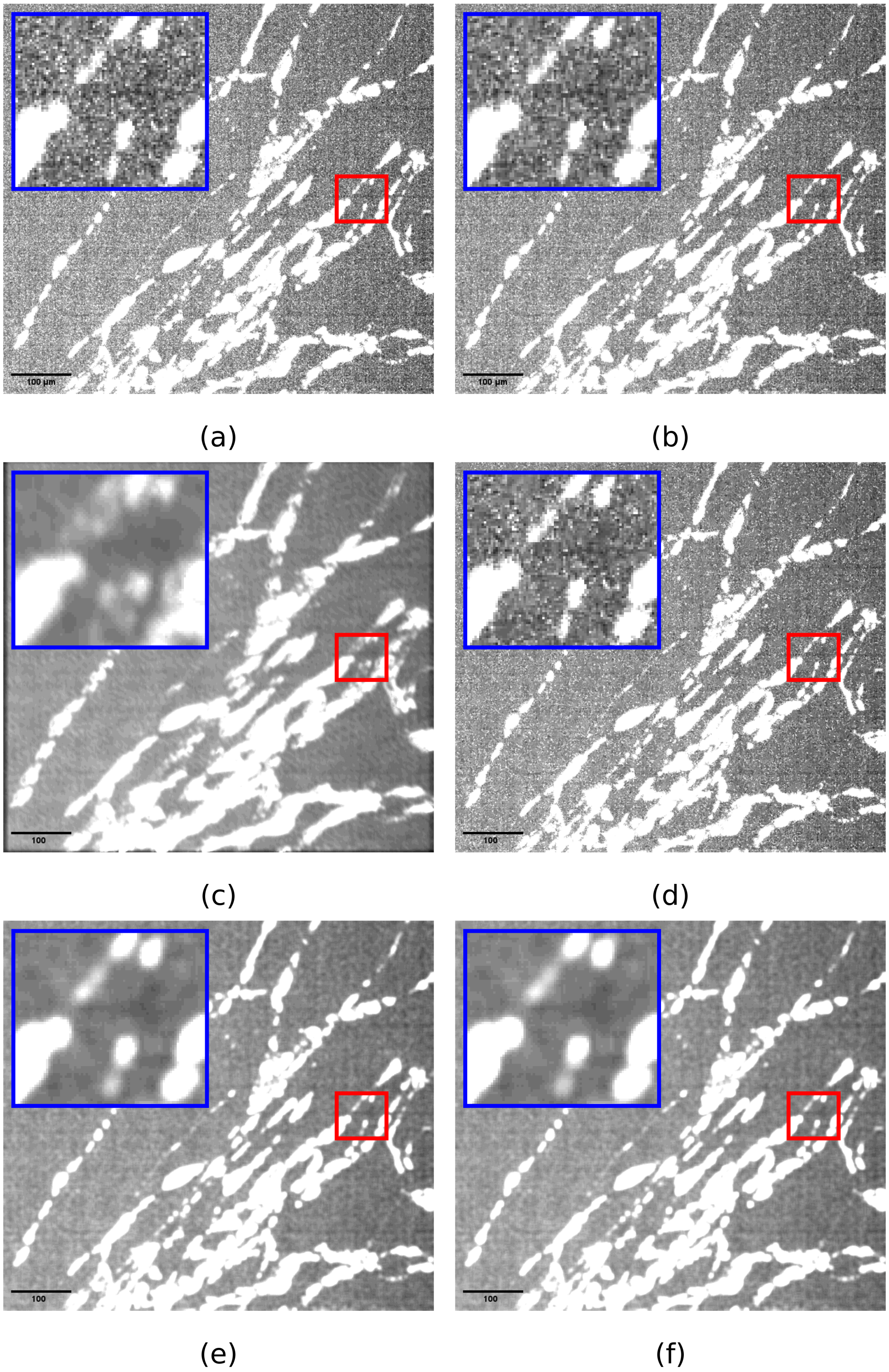


**Supplementary Figure S1.** Comparison of classical denoising filters applied to a representative MitoTracker fluorescence microscopy image. (a) Original raw fluorescence image containing genuine mitochondrial signal alongside speckle noise and isolated pixel artifacts in the background. (b-f) Results after applying Wavelet-Bayes (b), bilateral filtering (c), non-local means (d), Lee filter (e), and Kuan filter (f). Red squares indicate the region selected for magnified inspection; corresponding zoomed views are shown in blue squares. Scale bar: 100 μm.

Crucially, all of the above approaches operate on raw image intensities and therefore inevitably modify both signal and noise simultaneously, with no mechanism to distinguish topologically isolated artifacts from genuine biological structures of similar intensity. This fundamental limitation motivates the mask-based strategy of SPC-Clean, which foregoes intensity manipulation entirely and instead targets the topological structure of the thresholded foreground mask, enabling selective removal of isolated artifacts without any modification of the underlying fluorescence signal.

To further assess the limitations of classical filtering approaches, we applied the same five denoising methods: Wavelet-Bayes, bilateral filtering, non-local means (NLM), Lee, and Kuan, directly to the binary mask produced by global intensity thresholding, rather than to the raw fluorescence image (Figure S2).

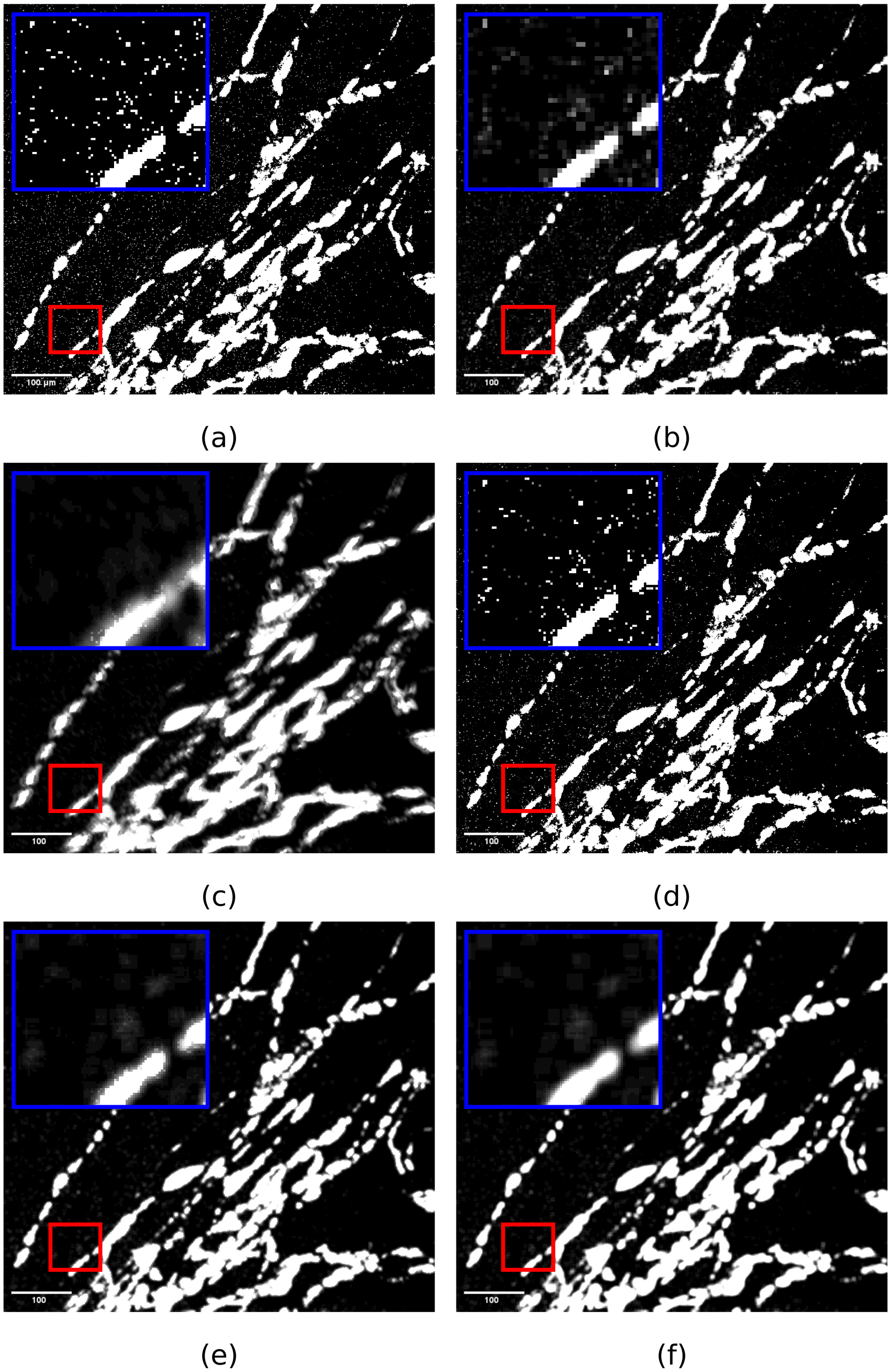


**Supplementary Figure S2.** Comparison of classical denoising filters applied to the binary mask after global intensity thresholding of a representative MitoTracker fluorescence microscopy image. (a) Binary mask generated by global thresholding of the raw fluorescence image, containing genuine mitochondrial foreground alongside isolated pixel artifacts and salt-and-pepper noise. (b–f) Results after applying Wavelet-Bayes (b), bilateral filtering (c), non-local means (d), Lee filter (e), and Kuan filter (f) to the thresholded binary mask. Red squares indicate the region selected for magnified inspection; corresponding zoomed views are shown in blue squares. Scale bar: 100 µm.

The results reveal distinct and complementary failure modes across the five methods. Wavelet-Bayes and non-local means, while designed for general-purpose noise suppression, fail to completely eliminate salt-and-pepper artifacts in the binary domain: residual isolated foreground pixels remain scattered across the background after filtering, indicating that these methods lack the spatial selectivity required to target sparse, disconnected noise clusters.

Bilateral, Lee, and Kuan filters achieve more aggressive background suppression, but at the cost of unacceptable signal distortion. Lee and Kuan filters, which assume a multiplicative speckle noise model, leave residual low-intensity shadow artifacts and diffuse background noise visible in the magnified regions, suggesting that their local homogeneity assumptions are violated by the sparse and heterogeneous foreground structure of thresholded mitochondrial masks. Bilateral filtering produces the most visually clean background but introduces the most severe signal distortion: spurious foreground regions appear that have no correspondence to genuine mitochondrial structures, and the boundaries of real foreground regions are substantially altered, expanding or merging adjacent structures in a manner that would directly corrupt downstream morphological measurements such as object size, shape, and connectivity.

Critically, all five methods share the same fundamental limitation: they operate on pixel intensity values and therefore cannot distinguish between an isolated noise pixel and a genuine low-intensity foreground pixel based on local intensity information alone. Any filter aggressive enough to suppress isolated noise will inevitably affect genuine signal of similar local intensity, and any filter conservative enough to preserve thin mitochondrial structures will leave residual noise intact. This intrinsic trade-off cannot be resolved by tuning filter parameters and motivates an entirely different approach. SPC-Clean sidesteps this limitation by operating on the topological connectivity of the binary mask rather than on pixel intensities, enabling selective removal of isolated artifacts based on their lack of neighborhood support rather than their intensity value, and leaving all retained foreground pixels completely unmodified.

To provide a direct comparison between speckle-specific filtering approaches and SPC-Clean, Figure S3 shows the results of applying the Lee filter, the Kuan filter, and SPC-Clean ($K = 3$) to the same thresholded binary mask derived from a representative MitoTracker fluorescence image.

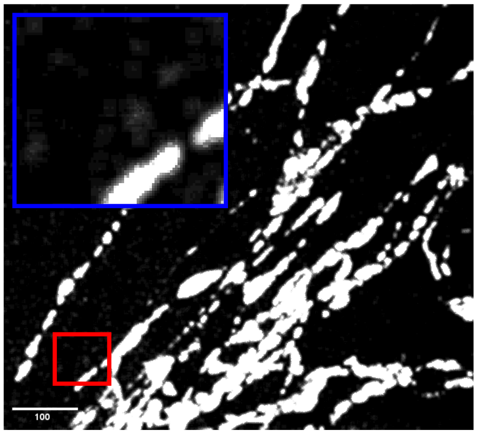


(a)

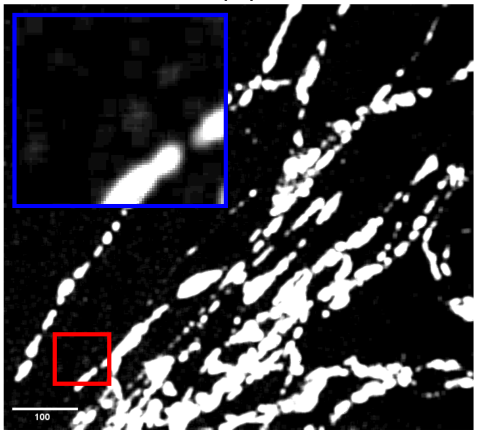


(b)

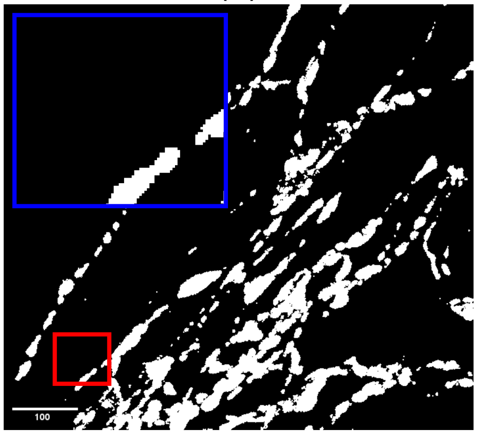


(c)

**Supplementary Figure S3.** Direct comparison of speckle-specific filters and SPC-Clean applied to a thresholded binary mask derived from a representative MitoTracker fluorescence microscopy image. (a) Result after applying the Lee filter to the binary mask. (b) Result after applying the Kuan filter to the binary mask. (c) Result after applying SPC-Clean ($K = 3$) to the binary mask. Red squares indicate the region selected for magnified inspection; corresponding zoomed views are shown in blue squares. Scale bar: 100 µm.

Lee and Kuan filters are among the most established methods for speckle noise suppression and are specifically designed to handle multiplicative noise under the assumption of locally homogeneous signal statistics. However, when applied to a binary mask, both filters exhibit a characteristic set of failure modes that render them unsuitable for this task. As visible in the magnified regions of Figure S3(a,b), neither filter completely eliminates isolated background pixels: residual noise remains scattered across the field of view, particularly in regions where noise pixels are clustered at a scale comparable to the filter's local window. More critically, both filters introduce low-intensity shadow artifacts in the background, diffuse foreground-like regions that do not correspond to any genuine biological structure, arising from the local averaging of sparse foreground pixels with their surrounding background. Furthermore, the boundaries of genuine mitochondrial structures are measurably altered: foreground regions are expanded, eroded, or merged with adjacent structures depending on local pixel density, directly corrupting morphological measurements such as object area, perimeter, and connectivity that are central to mitochondrial quantification pipelines.

SPC-Clean with $K = 3$ (Figure S3(c)) produces qualitatively and quantitatively superior results across all of these dimensions. Isolated foreground pixels and small spurious clusters are completely removed from the background, with no residual noise visible in the zoomed region. Crucially, the boundaries of genuine mitochondrial structures are preserved with high fidelity: no spurious signal is introduced, no existing foreground regions are merged or artificially expanded, and thin structural elements that would be vulnerable to boundary distortion by intensity-domain filters are retained intact. This behavior results directly from the design of SPC-Clean, which evaluates each foreground pixel solely on the basis of how many foreground neighbors it possesses: a pixel is removed only if it is topologically isolated, and retained unconditionally otherwise. Unlike Lee and Kuan filters, which modify all pixel values through local weighted averaging, SPC-Clean makes a binary keep-or-remove decision for each pixel independently, ensuring that retained pixels are identical to their original values and that no intermediate or shadow values are ever introduced into the mask.

Taken together, Figures S2 and S3 establish that speckle-specific intensity-domain filters are fundamentally mismatched to the problem of artifact suppression in binary fluorescence microscopy masks, and that SPC-Clean provides a more principled and effective solution by targeting topological isolation rather than local intensity statistics.

A key parameter governing the behavior of SPC-Clean is the minimum neighborhood support threshold $K$, which determines how many active foreground neighbors a pixel must possess in its 8-connected neighborhood in order to be retained in the mask. To characterize the effect of $K$ on cleaning performance and signal preservation, we evaluated SPC-Clean across four values ($K \in \{2, 3, 4, 5\}$) on a representative MitoTracker fluorescence image, with the intensity threshold fixed at $H = 128$ throughout (Figure S4).

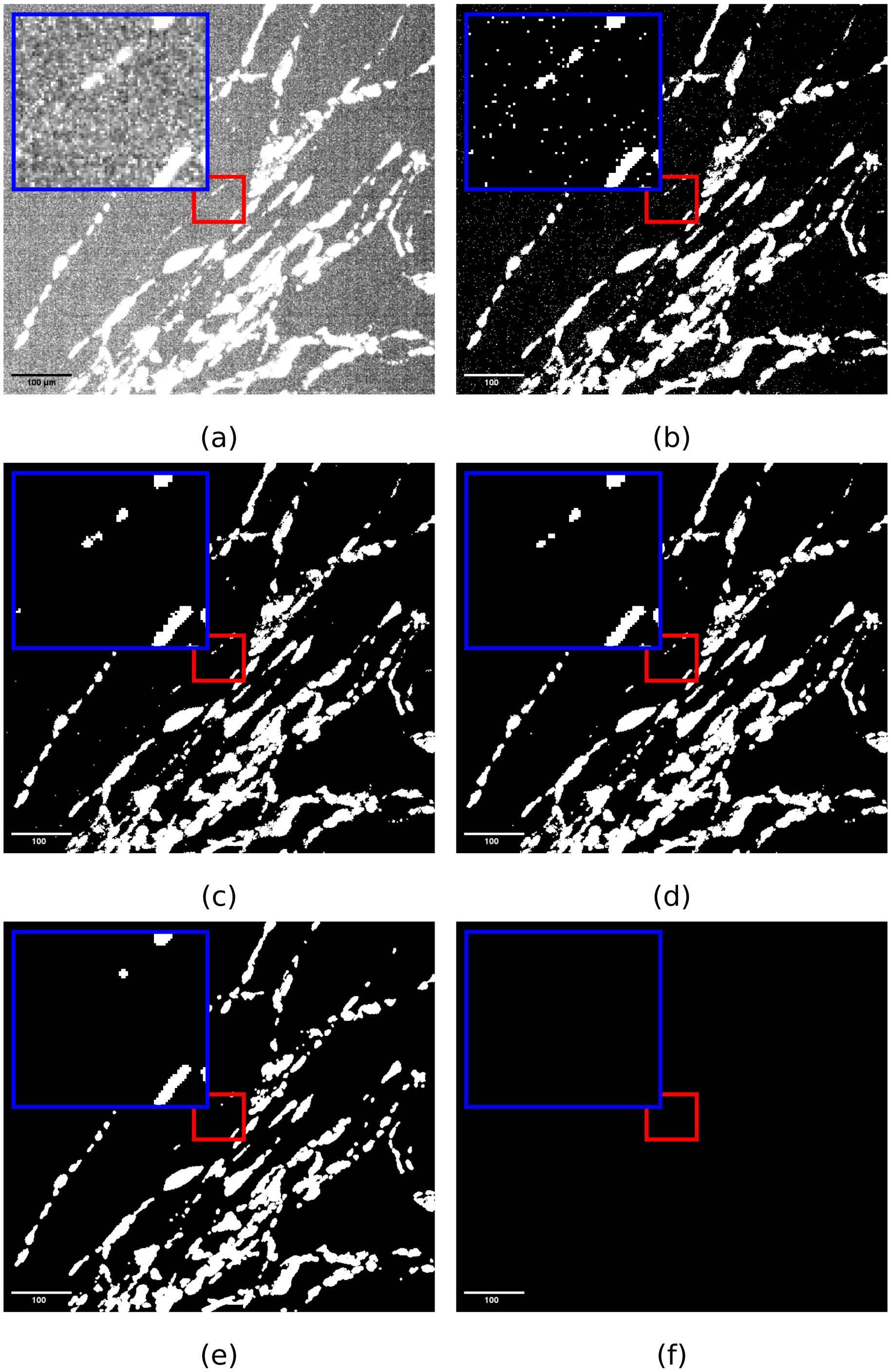


**Supplementary Figure S4.** Effect of the neighborhood support parameter $K$ on SPC-Clean output for a representative MitoTracker fluorescence microscopy image. (a) Original raw fluorescence image containing genuine mitochondrial signal alongside speckle noise and isolated pixel artifacts. (b) Binary mask generated by global thresholding at $H = 128$, revealing the salt-and-pepper noise introduced by thresholding. (c–f) SPC-Clean output masks for increasing values of the neighborhood support threshold: $K = 2$ (c), $K = 3$ (d), $K = 4$ (e), and $K = 5$ (f), all with $H = 128$. Red squares indicate the region selected for magnified inspection; corresponding zoomed views are shown in blue squares. Scale bar: 100 µm.

The original raw image (Figure S4(a)) contains genuine mitochondrial signal alongside background speckle noise and isolated pixel artifacts. Global thresholding at $H = 128$ (Figure S4(b)) produces a binary mask in which the mitochondrial network is captured but accompanied by a large number of salt-and-pepper noise pixels distributed across the background, illustrating the fundamental limitation of threshold-based segmentation.

At $K = 2$ (Figure S4(c)), SPC-Clean removes foreground pixels with fewer than two active neighbors, primarily targeting isolated pixels and very small foreground artifacts. Although this reduces some of the noise, small connected noise clusters can remain when their pixels have sufficient neighboring support. As a result, residual speckle artifacts are still visible in the background, indicating that $K = 2$ does not provide sufficient speckle reduction for this noise regime.

At $K = 3$ (Figure S4(d)), the neighborhood support requirement is sufficient to eliminate all isolated speckle artifacts and small spurious clusters, producing a clean binary mask in which the background is entirely free of noise pixels. Critically, genuine mitochondrial structures, including thin filamentous regions, are fully preserved at this setting, as they possess sufficient local connectivity to satisfy the $K = 3$ criterion throughout the iterative pruning process. We therefore identify $K = 3$ as the optimal operating point for MitoTracker fluorescence data and adopt this value for all quantitative experiments reported in this work.

At $K = 4$ (Figure S4(e)), the cleaning becomes over-aggressive: in addition to removing noise, the algorithm begins to suppress pixels at the tips and peripheries of genuine thin mitochondrial structures, which by their nature have fewer local neighbors than the interior of larger connected regions. This results in measurable erosion of real foreground signal and loss of fine structural detail that is biologically meaningful.

At $K = 5$ (Figure S4(f)), the neighborhood support requirement is so stringent that virtually no foreground pixel can satisfy it, and the algorithm removes essentially the entire foreground mask including the main mitochondrial network. The resulting mask is effectively empty and unsuitable for any downstream analysis.

Together, these results illustrate the characteristic behavior of SPC-Clean as a function of $K$: the method transitions from under-cleaning ($K = 2$) through an optimal operating regime ($K = 3$) to progressive signal erosion ($K = 4$) and complete foreground suppression ($K = 5$). The sharpness of this transition reflects the sparse and topologically thin nature of mitochondrial structures, where genuine foreground pixels typically have between three and seven active neighbors, placing the optimal $K$ squarely at the boundary between noise and signal in neighborhood-connectivity space. This analysis confirms that $K = 3$ provides a robust and interpretable operating point, and that the sensitivity of results to $K$ is predictable and monotonic, facilitating principled parameter selection in new datasets.

##### Parameter selection guidelines.

SPC-Clean exposes two parameters that should be selected based on the characteristics of the input fluorescence data: the intensity threshold $H$ and the neighborhood support threshold $K$.

The intensity threshold $H$ controls the initial foreground mask generated in Stage 1 and should be chosen to capture the genuine biological signal while minimizing gross background inclusion. In practice, $H$ can be determined using standard automated thresholding strategies adapted to the intensity distribution of the specific fluorescence channel: for images with a skewed, non-zero-baseline histogram characteristic of MitoTracker and similar organelle-specific dyes, the triangle threshold [Zack et al.(1977)] provides a principled and reproducible starting point. For images with a clearer bimodal intensity distribution, Otsu's method [Otsu(1979)] is appropriate. Importantly, the precise choice of $H$ has a limited effect on the final SPC-Clean output, as the subsequent neighborhood pruning step in Stage 2 is specifically designed to correct for the isolated noise pixels that any threshold will inevitably introduce. $H$ should therefore be set conservatively to avoid excluding genuine low-intensity signal, accepting that some noise will enter the initial mask and be removed by SPC-Clean in the following stage.

The neighborhood support threshold $K$ controls the aggressiveness of the topological pruning in Stage 2 and is the more sensitive of the two parameters. As demonstrated in Figure S4, the optimal value of $K$ depends on the connectivity characteristics of the biological structures of interest. For sparse, thin structures such as mitochondrial filaments, where genuine foreground pixels typically have three to seven active neighbors, $K = 3$ provides the optimal balance between complete noise suppression and full signal preservation. For denser or more compact structures, higher values of $K$ may be tolerated without signal loss. As a practical guideline, we recommend starting with $K = 3$ and inspecting the resulting mask on a representative image. If residual speckle remains, $K$ can be increased by one. If genuine thin structures begin to disappear, $K$ should instead be decreased. As shown in Figure S4, the effect of changing $K$ is monotonic and predictable. In practice, an appropriate value can therefore be selected by visually inspecting a small number of settings on a representative image, without requiring an extensive grid search or quantitative optimization. Once suitable values of $H$ and $K$ have been selected, they can be applied consistently across images acquired under the same experimental conditions. This approach was used for the 1,002 images from three independent acquisitions, for which SPC-Clean showed stable and reproducible performance, as reported in Supplementary Tables 2 and 3.

**Supplementary Table 2** Quantitative evaluation of SPC-Clean pooled across three MitoTracker fluorescence microscopy image fields of primary human fibroblasts acquired on an Opera Phenix spinning-disk confocal microscope ($N = 1{,}002$ images total; 334 image planes per field). Metrics were computed on raw fluorescence images using the initial threshold-derived foreground mask (*Before*) and the corresponding SPC-Clean-refined mask (*After*). Values are reported as mean ± std. Speckle components are defined as connected foreground regions with an area $\leq 10$ pixels. Lower speckle count and speckle fraction indicate greater speckle reduction. SSIM is computed between the raw-intensity-weighted pre-cleaning and post-cleaning foreground images within the bounding box containing the union of their foreground regions. Higher SSIM values indicate greater structural similarity between the pre-cleaning and post-cleaning foreground signal, with values $> 0.9$ indicating high structural similarity.

| **Metric** | **Before (mean ± std)** | **After (mean ± std)** | **Change** |
|---|---|---|---|
| Speckle count | $1113 \pm 1118$ | $35 \pm 73$ | −96.9% |
| Speckle fraction | $0.976 \pm 0.029$ | $0.291 \pm 0.308$ | −70.2% |
| SSIM | — | $0.992 \pm 0.005$ | — |

**Supplementary Table 3** Per-field quantitative evaluation of SPC-Clean across three MitoTracker fluorescence microscopy image fields of primary human fibroblasts acquired on an Opera Phenix spinning-disk confocal microscope (63× water objective, NA,=,1.15, 334-plane z-stacks, 0.3,µm axial step). All metrics are computed on raw fluorescence images. *Before*: initial threshold-derived foreground mask applied to the raw image. *After*: corresponding SPC-Clean-refined mask. Values are reported as mean ± std over all image planes within each field. Speckle components are defined as connected foreground regions with an area ≤ 10 pixels. Lower speckle count and speckle fraction indicate greater speckle reduction. SSIM is computed between the raw-intensity-weighted pre-cleaning and post-cleaning foreground images within the bounding box containing the union of their foreground regions. Higher SSIM values indicate greater structural similarity between the pre-cleaning and post-cleaning foreground signal, with values > 0.9 indicating high structural similarity.

| | **Run 1** | | **Run 2** | | **Run 3** | |
|---|---|---|---|---|---|---|
| **Metric** | **Before** | **After** | **Before** | **After** | **Before** | **After** |
| Speckle count | $582 \pm 152$ | $6 \pm 8$ | $1744 \pm 1614$ | $71 \pm 110$ | $1014 \pm 662$ | $27 \pm 40$ |
| Speckle count change (%) | −99.0% | | −95.9% | | −97.4% | |
| Speckle fraction | $0.976 \pm 0.032$ | $0.199 \pm 0.273$ | $0.977 \pm 0.027$ | $0.357 \pm 0.332$ | $0.974 \pm 0.029$ | $0.317 \pm 0.297$ |
| SSIM | $0.995 \pm 0.001$ | | $0.990 \pm 0.006$ | | $0.991 \pm 0.005$ | |

### S5. Additional Experimental Results

To assess the generalizability of SPC-Clean beyond the primary validation dataset, we applied the method to three additional independent fluorescence microscopy datasets representing distinct biological samples, labeling strategies, and imaging conditions. In all cases, SPC-Clean was applied identically, with no parameter adjustment relative to the primary experiments, demonstrating that the method is robust across diverse fluorescence microscopy contexts.

##### MitoTracker-stained HeLa cells (Supplementary Figure S5)

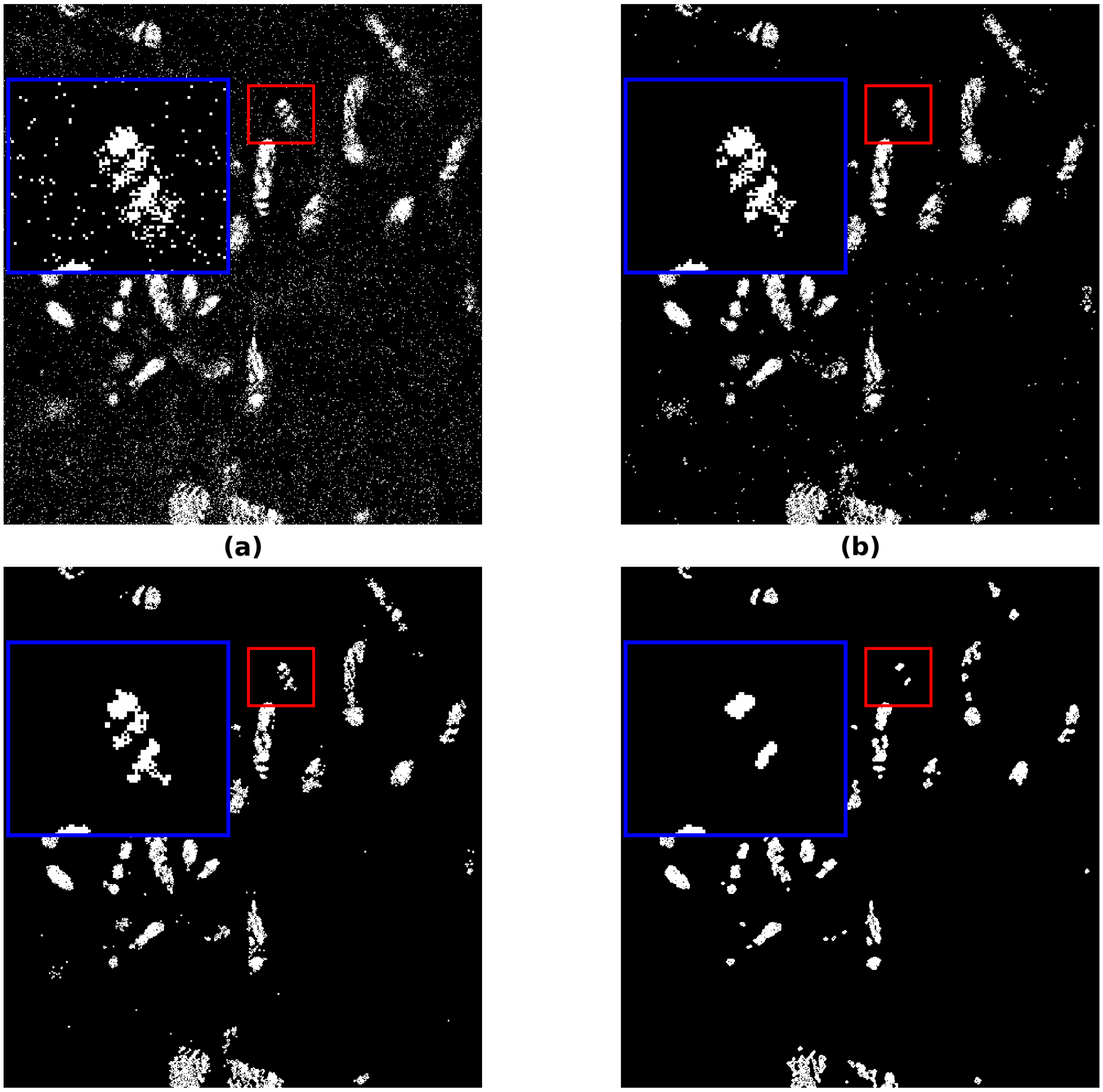

(a) (b) (c) (d)

**Supplementary Figure S5.** SPC-Clean applied to a MitoTracker-stained HeLa cell image, demonstrating the effect of the neighborhood support parameter $K$ on speckle reduction and structural preservation. (a) Binary mask generated by global thresholding at $H = 118$, revealing extensive salt-and-pepper noise and isolated foreground artifacts introduced by thresholding alongside genuine mitochondrial signal. (b–d) SPC-Clean output masks with $K = 2$ (b), $K = 3$ (c), and $K = 4$ (d), all with $H = 118$. Red squares indicate the region selected for magnified inspection; corresponding zoomed views are shown in blue squares.

Global thresholding of the raw MitoTracker channel at $H = 118$ produced a binary mask containing extensive salt-and-pepper noise and numerous isolated foreground pixels distributed across the background, as shown in Supplementary Figure S5(a). Despite the high image quality achieved by pan-ExM, thresholding alone is insufficient to produce a clean foreground mask, confirming that speckle artifacts arise independently of imaging modality and sample preparation protocol.

SPC-Clean was applied to this mask across three values of the neighborhood support parameter $K$. At $K = 2$ (Supplementary Figure S5(b)), the majority of isolated noise pixels are removed, producing a substantially cleaner mask than the raw threshold output. However, residual artifacts remain visible in the background, particularly in the zoomed region, where small noise clusters with at least two mutually neighboring pixels survive the $K = 2$ pruning criterion. At $K = 3$ (Supplementary Figure S5(c)), all isolated speckle artifacts and spurious foreground clusters are completely eliminated. The connected mitochondrial network is fully preserved, including thin tubular structures characteristic of HeLa mitochondria, demonstrating that genuine foreground pixels possess sufficient neighborhood connectivity to satisfy the $K = 3$ criterion throughout the iterative pruning process. At $K = 4$ (Supplementary Figure S5(d)), the neighborhood support requirement becomes over-aggressive for this dataset: pixels at the tips and peripheries of thin mitochondrial filaments, which by their nature have fewer active neighbors than interior regions, fail to meet the $K = 4$ criterion and are incorrectly removed. This results in measurable erosion and fragmentation of genuine mitochondrial structures, producing a mask that underrepresents the true mitochondrial network and would introduce systematic errors into downstream morphological quantification. These results are fully consistent with the parameter sensitivity analysis reported for the primary dataset (Figure S4) and confirm that $K = 3$ represents the optimal operating point across independent datasets and imaging conditions.

**Antibody-labeled HeLa cells. (Supplementary Figure S6).**

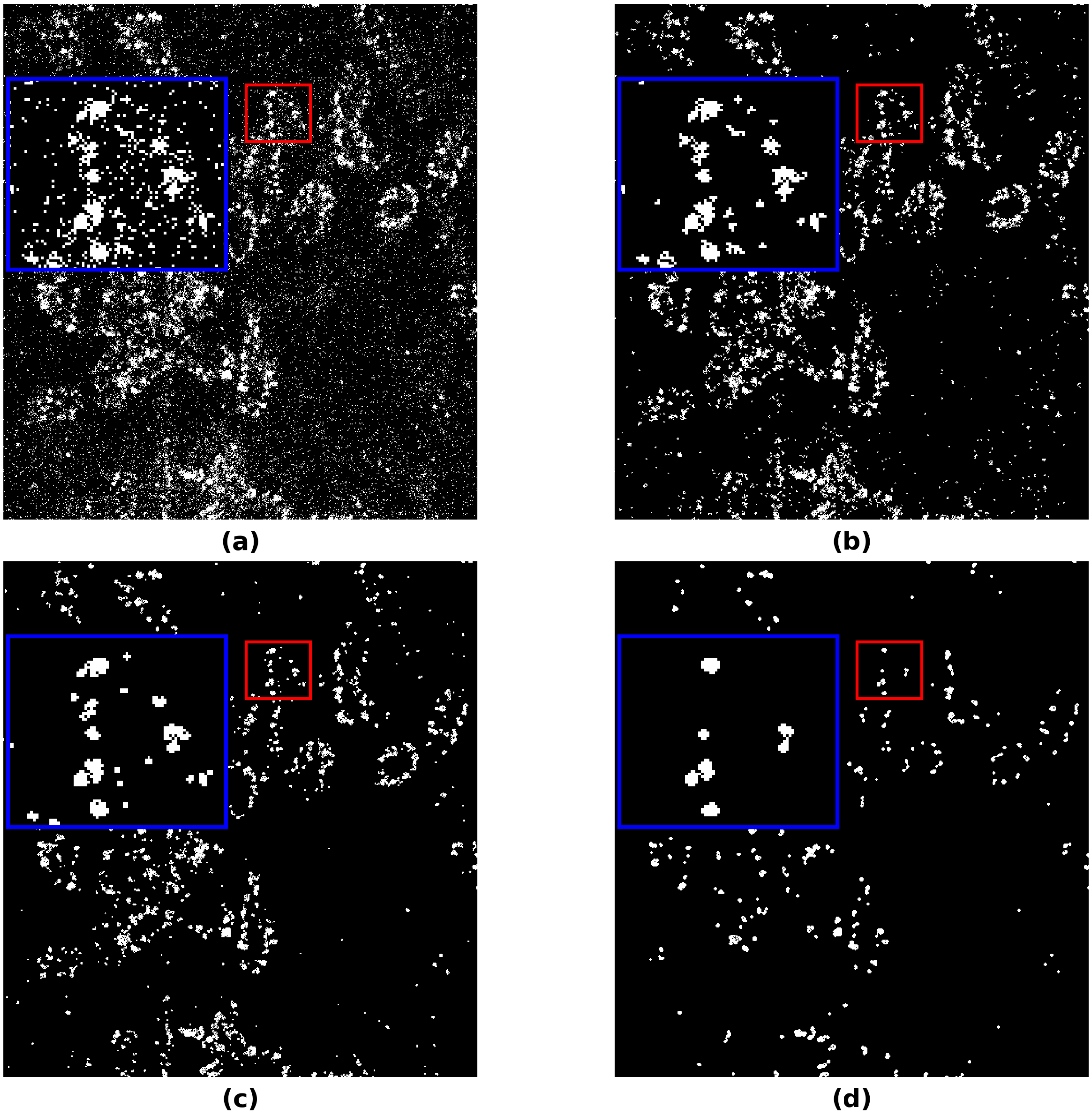

**Supplementary Figure S6.** SPC-Clean applied to an antibody-labeled HeLa cell image processed with pan-Expansion Microscopy (pan-ExM), demonstrating the effect of the neighborhood support parameter $K$ on noise suppression and signal preservation in the immunofluorescence channel. (a) Binary mask generated by global thresholding at $H = 118$, revealing dense salt-and-pepper noise and isolated foreground artifacts arising from diffuse background fluorescence and nonspecific antibody labeling. (b–d) SPC-Clean output masks with $K = 2$ (b), $K = 3$ (c), and $K = 4$ (d), all with $H = 118$. Red squares indicate the region selected for magnified inspection; corresponding zoomed views are shown in blue squares.

The same HeLa cell preparation described above was additionally labeled using indirect immunofluorescence within the pan-ExM framework [Tian et al.(2025)]. The antibody-labeled images show a different and more challenging noise profile than the MitoTracker-stained images. After thresholding, the resulting masks contain more diffuse background signal and a higher number of isolated foreground artifacts, as shown in Supplementary Figure S6(a). The resulting binary mask at $H = 118$ contains substantially more spurious foreground pixels than the MitoTracker channel, reflecting the inherently lower specificity of indirect immunofluorescence relative to organelle-targeted dyes. This makes the antibody channel a more demanding test case for SPC-Clean, as the higher noise density increases the risk of residual artifacts surviving the pruning step.

At $K = 2$ (Supplementary Figure S6(b)), SPC-Clean suppresses the majority of isolated noise pixels, producing a noticeably cleaner mask. However, consistent with the observations from the MitoTracker dataset, residual artifacts remain in the background, particularly in regions where noise pixels form small locally connected clusters that satisfy the $K = 2$ neighborhood criterion. At $K = 3$ (Supplementary Figure S6(c)), all spurious foreground pixels and isolated noise clusters are completely eliminated across the entire field of view. Genuine immunolabeled structures are fully preserved, including fine structural details visible in the zoomed region that would be vulnerable to distortion or loss by conventional intensity-domain filtering approaches. The clean separation between noise and signal achieved at $K = 3$ confirms that immunolabeled structures, despite their heterogeneous intensity distribution and irregular morphology, possess sufficient local neighborhood connectivity to consistently satisfy the pruning criterion. At $K = 4$ (Supplementary Figure S6(d)), over-aggressive pruning again becomes apparent: genuine immunolabeled structures begin to be eroded at their boundaries and peripheries, with thin or weakly connected regions incorrectly classified as insufficiently supported and removed from the mask. This would introduce systematic underestimation of labeled structure size and density in downstream quantification.

Taken together, the results from the MitoTracker and antibody channels of HeLa cells demonstrate that $K = 3$ generalizes across two distinct fluorescence labeling strategies within the same pan-ExM experimental framework, and that the optimal operating point is consistent across channels with substantially different noise profiles and foreground densities.

**Antibody-labeled mouse kidney tissue sections (Supplementary Figure S7).**

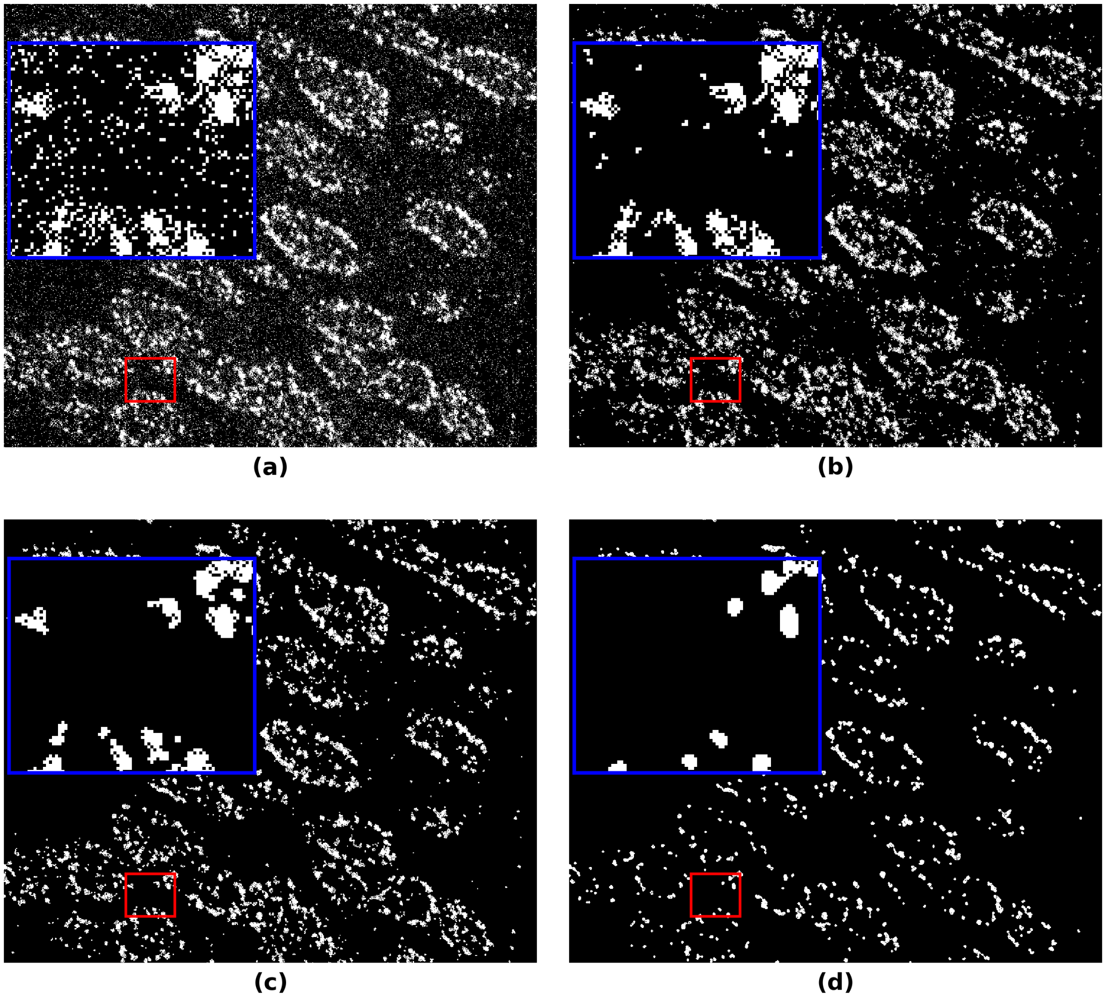


**Supplementary Figure S7.** SPC-Clean applied to an antibody-labeled mouse kidney tissue section processed with pan-Expansion Microscopy for tissue (pan-ExM-t), demonstrating the effect of the neighborhood support parameter $K$ on noise suppression and signal preservation in a complex tissue context. (a) Binary mask generated by global thresholding at $H = 110$, revealing extensive salt-and-pepper noise and isolated foreground artifacts arising from autofluorescence, nonspecific antibody labeling, and uneven labeling density characteristic of tissue sections. (b–d) SPC-Clean output masks with $K = 2$ (b), $K = 3$ (c), and $K = 4$ (d), all with $H = 110$. Red squares indicate the region selected for magnified inspection; corresponding zoomed views are shown in blue squares.

To evaluate SPC-Clean on tissue rather than cultured cells, we applied the method to fluorescence images of mouse kidney sections prepared using the pan-ExM-t protocol optimized for tissue expansion [Tian et al.(2025)].

Compared with the cultured HeLa cell images, the kidney tissue images showed a more heterogeneous fluorescence intensity distribution, making foreground separation with a single global threshold more challenging. While $H = 118$ provided an appropriate initial foreground mask for the HeLa cell datasets, a lower threshold of $H = 110$ was used for the kidney tissue dataset to retain the relevant immunolabeled structures. This difference illustrates why $H$ should be selected according to the intensity characteristics of the dataset rather than fixed across different sample types. A lower threshold can also retain more isolated foreground artifacts in the initial mask. The subsequent SPC-Clean pruning step in Stage 2 addresses these artifacts by removing pixels with insufficient local neighborhood support while retaining more spatially coherent foreground structures.

As shown in Supplementary Figure S7(a), global thresholding at $H = 110$ produces a binary mask with dense and spatially irregular salt-and-pepper noise across the background. The density of these noise pixels varies considerably between regions with different levels of autofluorescence. At the same time, the kidney tissue contains complex biological structures, including thin tubular elements and glomerular regions with heterogeneous foreground density. The combination of these structures with spatially variable background noise makes the kidney tissue dataset the most challenging of the three supplementary datasets evaluated with SPC-Clean.

At $K = 2$ (Supplementary Figure S7(b)), SPC-Clean removes a substantial fraction of isolated noise pixels. However, residual artifacts persist in regions of elevated autofluorescence, where locally clustered noise pixels satisfy the $K = 2$ neighborhood criterion and therefore survive the pruning step. This behavior confirms that $K = 2$ is systematically insufficient for complete noise suppression in tissue samples, where the higher noise density increases the probability of isolated noise pixels forming small locally connected clusters.

At $K = 3$ (Supplementary Figure S7(c)), SPC-Clean produces a substantially cleaner binary mask, with isolated speckle artifacts and small spurious foreground clusters strongly reduced across the field of view. This improvement is also evident in regions with higher autofluorescence, where residual artifacts remain at $K = 2$. The effectiveness of $K = 3$ in this more complex tissue environment can be explained by the local topology of the foreground mask. Speckle artifacts and isolated noise pixels typically occur as single pixels or small, weakly connected clusters and therefore have relatively few active neighbors. In contrast, genuine tissue structures generally form larger and more spatially connected foreground regions with greater local neighborhood support. This difference in local connectivity allows SPC-Clean to preferentially remove small foreground artifacts while retaining the major structural features of the tissue. In this dataset, $K = 3$ therefore provided a suitable balance between speckle reduction and structural retention without requiring adjustment from the value used for the cell-culture datasets. Thin tubular elements, glomerular boundaries, and spatially heterogeneous foreground regions remain visible and connected in the cleaned mask, as illustrated by the zoomed region in Supplementary Figure S7(c).

At $K = 4$ (Supplementary Figure S7(d)), the effect of more aggressive pruning becomes apparent. Thin tubular structures and peripheral foreground pixels along glomerular boundaries may no longer satisfy the $K = 4$ neighborhood criterion and are consequently removed. This leads to visible fragmentation of some continuous tubular structures and loss of fine structural detail. Preserving these features is particularly important in kidney pathology, where tubular morphology is used to assess changes such as tubular dilation, atrophy, and cross-sectional area, including in models of acute kidney injury such as the cisplatin model evaluated here. Excessive removal of these structural features could therefore affect subsequent quantitative morphological analyses.

These results highlight an important practical distinction between the two SPC-Clean parameters. The intensity threshold $H$ is sample-dependent and must be adjusted to reflect the background intensity characteristics of the specific biological preparation: lower values are appropriate for tissue samples with elevated autofluorescence, while higher values suit cleaner cell culture preparations. The neighborhood support threshold $K$, by contrast, is governed by the universal topological distinction between speckle noise and genuine biological signal: noise pixels are sparsely connected regardless of sample type, while real structures are spatially coherent. This explains why $K = 3$

provided a consistent and effective balance between speckle reduction and structural retention across all three datasets, including MitoTracker-stained HeLa cells, antibody-labeled HeLa cells, and antibody-labeled mouse kidney tissue sections. Despite differences in sample type, imaging conditions, labeling strategies, and noise characteristics, the same value of $K = 3$ was applied successfully without dataset-specific adjustment.

##### Summary.

Across the supplementary datasets, SPC-Clean was evaluated on two different biological sample types, cultured HeLa cells and mouse kidney tissue sections, using two fluorescence-labeling strategies, organelle-specific MitoTracker labeling and indirect immunofluorescence, as well as two sample-preparation protocols, cell-culture pan-ExM and tissue pan-ExM-t. Across these different conditions, $K = 3$ consistently provided strong reduction of isolated speckle artifacts and salt-and-pepper noise in the threshold-derived binary masks while retaining the major biological structures.

Notably, the same neighborhood parameter $K = 3$ was used across all supplementary datasets without dataset-specific adjustment. This consistent performance can be explained by the local topology targeted by SPC-Clean. Isolated noise pixels and small speckle clusters generally have few active neighbors, whereas biological structures such as mitochondrial filaments in cultured cells and tubular or glomerular structures in kidney tissue tend to form more spatially connected foreground regions. SPC-Clean uses this difference in local connectivity to preferentially remove small, weakly connected foreground components while retaining more coherent structures.

In contrast to $K$, the intensity threshold $H$ was adjusted between datasets to account for differences in fluorescence intensity and background levels, in accordance with the parameter-selection guidelines described above. Taken together, these results show that SPC-Clean can be applied across different biological samples, including tissue sections, fluorescence-labeling strategies, and sample-preparation protocols. They also demonstrate that the same $K = 3$ setting can provide a consistent balance between speckle reduction and structural retention across substantially different experimental conditions.
